# Tandem Mass Tag-Based High-Resolution LC-MS/MS identifies free D-aspartate-induced expression of proteins linked to schizophrenia and autism spectrum disorder

**DOI:** 10.1101/2025.04.18.649487

**Authors:** Francesco Errico, Rosita Russo, Federica Carrillo, Tommaso Nuzzo, Raffaella di Vito, Enza Canonico, Paolo Vincenzo Pedone, Ferdinando Di Cunto, Teresa Esposito, Alessandro Usiello, Angela Chambery

**Author notes:** These authors contributed equally to this work (Francesco Errico, Rosita Russo). **Corresponding authors:** Angela Chambery: Department of Environmental, Biological and Pharmaceutical Science and Technologies, University of Campania “Luigi Vanvitelli”, 81100 Caserta, Italy;., Alessandro Usiello: Laboratory of Translational Neuroscience, CEINGE Biotecnologie Avanzate “Franco Salvatore”, 80145 Naples, Italy; Department of Environmental, Biological and Pharmaceutical Science and Technologies, University of Campania “Luigi Vanvitelli”, 81100 Caserta, Italy;.

## Abstract

D-aspartate is an endogenous agonist of NMDA and mGlu5 receptors, with a distinctive spatiotemporal expression profile that peaks in the prenatal and early postnatal brain. This suggests a critical role for D-aspartate metabolism in modulating neurodevelopmental processes linked to glutamatergic neurotransmission. However, the precise mechanisms through which D-aspartate exerts its effects remain unclear. To elucidate the molecular pathways orchestrated by early D-aspartate signalling, we employed a knockin mouse model characterized by constitutive D-aspartate depletion due to the prenatal expression of its degradative enzyme, D-aspartate oxidase. Using an advanced quantitative proteomic approach based on Tandem Mass Tag isobaric labelling and nano-liquid chromatography coupled with high-resolution tandem mass spectrometry, we investigated the proteomic variations induced by D-aspartate depletion during postnatal brain development comparing *Ddo* knockin mice with their wild-type littermates. Our findings reveal that D-aspartate modulates the neonatal expression of proteins involved in glutamatergic neurotransmission, nervous system development, and cytoskeleton organization. Moreover, proteomic analysis identified a subset of D-aspartate-regulated proteins mapping molecular pathways associated with autism spectrum disorder and schizophrenia. These findings offer new perspectives on the complex protein networks influenced by D-aspartate metabolism in the developing brain and highlight its potential impact on cerebral function in health and psychiatric disorders.

## Introduction

Free D-aspartate (D-Asp) is selectively present in high amounts in the mammalian brain during prenatal and early postnatal development^1–3^. The distinctive spatiotemporal profile of cerebral D-Asp occurrence is tightly regulated by the postnatal expression of D-aspartate oxidase (DDO)^4–6^, the enzyme that catalyzes its selective degradation^7, 8^. While the degrading mechanism of D-Asp is well-established, its biosynthesis in mammals remains unclear. However, studies in mice suggest that serine racemase (SR), the enzyme responsible for D-serine biosynthesis^9^, may also function as a D-Asp synthase in the forebrain due to its ability to catalyze the chiral conversion of L-aspartate^10, 11^.

Pharmacological studies support the idea that D-Asp acts as an endogenous agonist at the glutamate GluN2 site of ionotropic NMDA receptors (NMDARs)^2, 12–15^ and at metabotropic Glu5 receptors (mGluR5)^16^. Microdialysis experiments in adult freely moving mice have further demonstrated that D-Asp crosses the blood-brain barrier and is present at nanomolar concentrations in the extracellular space of the cortex^6, 17^. Moreover, pharmacological oral and intraperitoneal administration of D-Asp in adult mice triggers a significant *in vivo* release of L-glutamate in the prefrontal cortex (PFC) of freely moving mice^17^, providing further evidence of its modulatory role on glutamatergic neurotransmission.

In line with its neuroactive role, studies in rodents and humans have shown that upregulation of D-Asp levels in *Ddo* knockout or D-Asp-treated mice stimulates brain activity, synaptic transmission and plasticity^15, 18–23^, and increases dendritic growth and spine density in the PFC and hippocampus^20, 24^. Specifically, in these brain regions, increased D-Asp levels appear to regulate both functional and structural synaptic plasticity through actin polymerization mechanisms^20, 24^.

To further explore the role of this atypical amino acid in regulating the downstream events of NMDAR and mGluR5 activation during brain development, we generated a knockin mouse model with prenatal and postnatal depletion of D-Asp, achieved through constitutive expression of the *Ddo* gene (*Ddo*-KI)^5^. Neuroanatomical investigations revealed that D-Asp depletion significantly perturbs brain morphology, with long-lasting effects into adulthood, as shown by the altered number of parvalbumin-positive GABAergic interneurons in the cortex of *Ddo*-KI mice^5^. Additionally, bromodeoxyuridine (BrdU) incorporation and immunohistochemical experiments showed a lower number of proliferating cells in the dorsal pallium during corticogenesis in these mice, which is reflected in a reduced grey matter volume in the adult cortex and striatum, as measured by structural magnetic resonance imaging (MRI)^25^.

Although the data presented above underscores the role of D-Asp in modulating glutamatergic transmission and neurodevelopment, its specific influence in regulating protein expression downstream NMDAR and mGluR5 stimulation in the developing brain remains unknown. In the present study, we sought to investigate the consequences of D-Asp deprivation at the proteome level in the brain of *Ddo*-KI mice at early postnatal stages using an advanced quantitative proteomic approach.

## Results

### Constitutive overexpression of *Ddo* gene leads to stable and specific depletion of D-aspartate levels in the *Ddo* knockin mouse brain

First, we confirmed *Ddo* gene overexpression in the whole brain of *Ddo*-KI mice at early postnatal stages, such as postnatal day 3 (P3) and P14, as well as at mature adult stage (around P300), compared to age-matched *Ddo* wild-type littermates (*Ddo*-WT). Quantitative real-time PCR (qRT-PCR) showed a significant increase in *Ddo* mRNA expression at each age analyzed (*Ddo*-KI vs *Ddo*-WT: P3, *p* = 0.0002; P14, *p* = 0.0355; P300, *p* < 0.001; Student’s *t*-test; Suppl. Fig. 1).

Using the same cohort of mice, we then evaluated the impact of constitutive *Ddo* overexpression on cerebral levels of D-Asp, its precursor, L-aspartate, and other amino acids directly or indirectly involved in glutamatergic transmission, including L-glutamate, L-glutamine, D-serine, L-serine, glycine, gamma-aminobutyric acid (GABA), taurine, L-asparagine, L-arginine, L-alanine and L-threonine. Additionally, we analyzed key ratios, such as D-Asp/total Asp (%), D-serine/total serine (%), glycine/L-serine, L-glutamine/L-glutamate and GABA/L-glutamate to assess potential alterations in amino acid metabolic interconversion. Statistical analysis revealed that *Ddo* overexpression selectively alters D-Asp levels (two-way ANOVA: genotype, F_(1,28)_ = 407.1 *p* < 0.0001; genotype x age F_(2,28)_ = 49.28, *p* < 0.0001) and D-Asp/total Asp ratio (genotype, F_(1,28)_ = 427.0, *p* < 0.0001; genotype x age F_(2,28)_ = 40.13, *p* < 0.0001) leading to D-Asp depletion at all analyzed ages (D-Asp and D-Asp/total Asp, *p* < 0.0001, Fisher’s *post-hoc* analysis) (Fig. 1a,c). In contrast, the levels of L-Asp and all other amino acids and metabolic ratios remained unchanged between genotypes (Fig. 1b,d-r).

**Figure 1.**
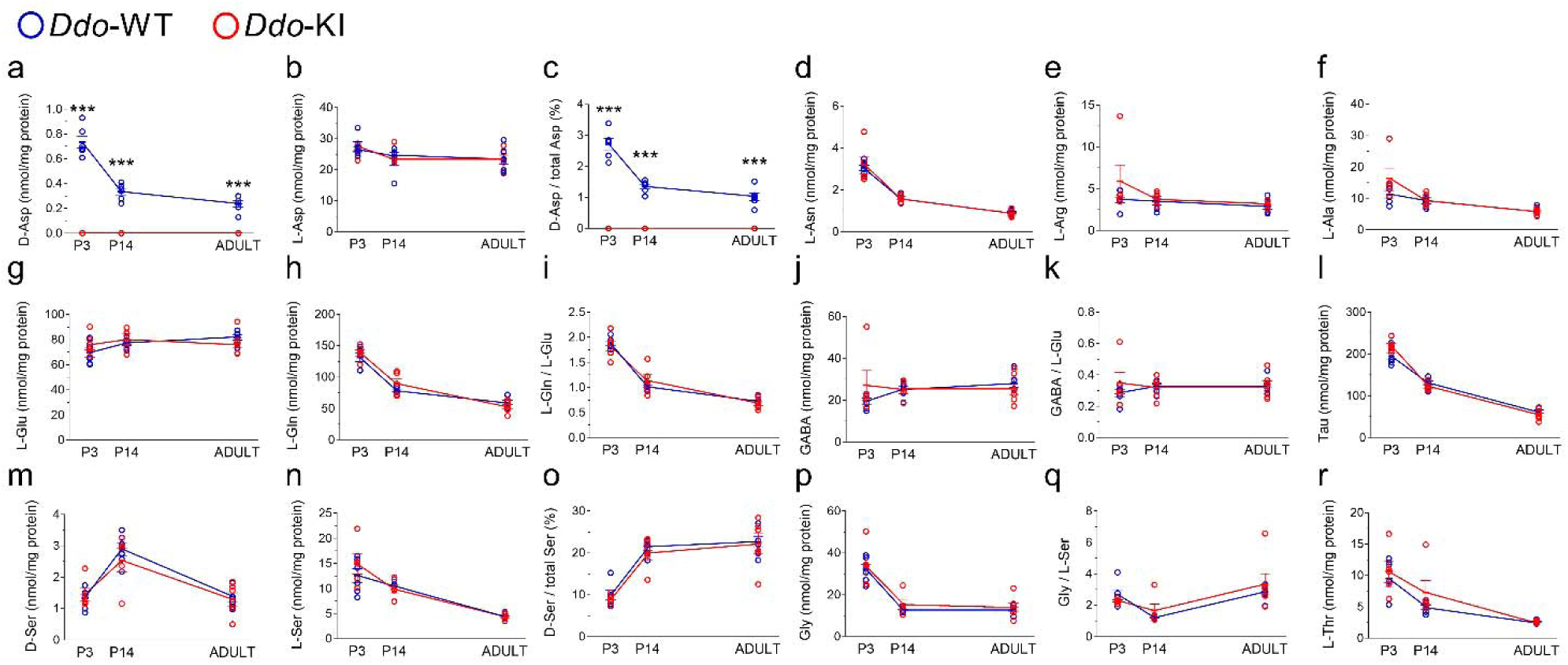
Amino acid levels in the whole brain of Ddo-KI mice during lifetime. Analysis of **(a)** D-aspartate and **(b)** L-aspartate levels, **(c)** D-aspartate/total aspartate ratio, **(d)** L-asparagine, **(e)** L-arginine, **(f)** L-alanine, **(g)** L-glutamate and **(h)** L-glutamine levels, **(i)** L-glutamine/L-glutamate ratio, **(j)** GABA levels, **(k)** GABA/L-glutamate ratio, **(l)** taurine, **(m)** D-serine and **(n)** L-serine levels, **(o)** D-serine/total serine ratio, **(p)** glycine levels, **(q)** glycine/L-serine ratio, and **(r)** L-threonine levels in the whole brain of *Ddo*-KI and *Ddo*-WT mice at post-natal day 3 (P3), P14 and P300. The amino acid content was expressed as nmol/mg protein, while the ratios were expressed as % (D-aspartate/total aspartate and D-serine/total serine) or absolute values (L-glutamine/L-glutamate, GABA/L-glutamate and glycine/L-serine). In each sample, free amino acids were detected in a single run. \*\*\**p* < 0.0001, compared with age-matched *Ddo*-WT mice (Fisher’s *post-hoc* comparison). Dots represent the single subject’s values while bars illustrate the median with interquartile range.

### A quantitative proteomics approach reveals that D-aspartate depletion affects proteome expression in early postnatal mouse brain

To investigate protein expression changes in *Ddo*-KI mice compared to the *Ddo*-WT littermates at different developmental stages, we applied a quantitative proteomic approach based on tandem mass tag (TMT) isobaric labelling and nano-liquid chromatography coupled with high-resolution tandem mass spectrometry (LC-MS/MS) (Fig. 2a). Specifically, we analyzed whole brains from *Ddo*-KI and *Ddo*-WT mice at P3, P7 and P14. We used an isobaric labelling-based workflow combined with LC-MS/MS on a Q-Exactive mass spectrometer^26^. A high number of peptide groups (24,716) was used for protein identification, with more than 80% used as unique peptides for protein quantification, confirming the high efficiency of peptide labelling (data not shown).

**Figure 2.**
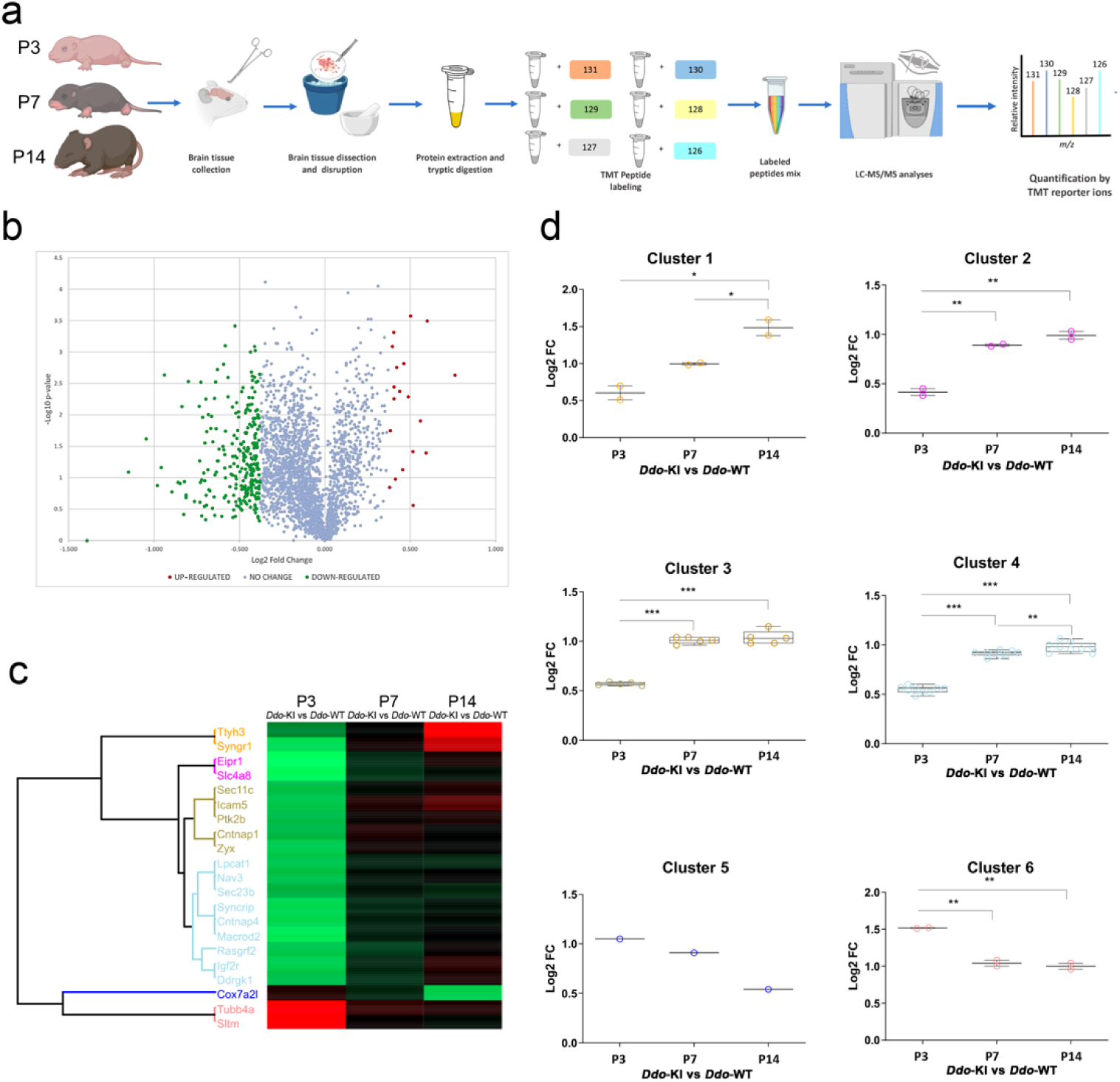
Quantitative proteomic approach reveals differentially expressed proteins in the whole brain of post-natal day 3 Ddo-KI mice. **(a)** Experimental design and proteomic workflow for comprehensive quantitative analysis of whole brains from *Ddo*-KI and *Ddo*-WT mice collected at different post-natal developmental stages, including postnatal day 3 (P3), P7 and P14. For Tandem Mass Tag (TMT) isobaric labelling, proteins extracted from whole brains were digested into peptides and labelled with TMT isobaric stable isotope tags. After sample mixing, peptides were analysed by LC-ESI-MS/MS. In MS1, the peptides appear as a single precursor. When fragmented during MS2, in addition to the normal fragment ions, the reporter regions dissociated to produce ion signals, which provided accurate quantitative information regarding the relative amount of the peptide in the samples. Image created with BioRender.com (www.biorender.com). **(b)** Volcano plot obtained from TMT-based quantitative proteomics analysis of P3 *Ddo*-KI versus age-matched *Ddo*-WT brain samples. Each point represents the difference in expression (Log2 fold-change) between the *Ddo*-KI and *Ddo*-WT plotted against the - Log10 p-value. Identified proteins with no changes in their regulation level are reported in light blue. Up- and down-regulated proteins (-0.4 ≥ Log2 fold-change ≥ 0.4) are indicated in red and green, respectively. **(c)** Hierarchical clustering (Euclidean average distance) of normalized protein intensities for differentially expressed proteins -0.6 ≥ (log2fold change) ≥ 0.6 identified in whole brains from *Ddo*-KI and *Ddo*-WT mice collected at postnatal days P3, P7 and P14 by quantitative nanoLC MS/MS. Down-regulated and up-regulated proteins in the heatmap are coloured in green and red, respectively. **(d)** Protein expression profiles of each cluster.

MS/MS and database search identified and quantified 2,594 non-redundant proteins, each with more than one peptide in at least two out of three replicates (Suppl. Data 1). Approximately 60% of proteins were commonly identified, while only about 21% of proteins were found to be uniquely identified. We extracted subsets of differentially expressed proteins from this dataset, applying a 1.3-fold change threshold, in at least one of the three postnatal comparisons (Suppl. Data 2). In detail, most of the differentially expressed proteins (288 out of 295) were observed at P3 stage in *Ddo*-KI samples, as illustrated in the volcano plot reporting -Log10 p-values versus Log2 fold-change values of *Ddo*-KI samples, compared with *Ddo*-WT (Fig. 2b and Suppl. Data 2). Following data filtering, we identified a subset of twenty-one proteins whose relative expression levels changed 1.5-fold or more in at least one time-point (P3, P7, P14) compared to *Ddo*-WT samples (Table 1; Fig. 2c). Proteomic analysis indicated that in four out of six clusters (clusters 1-4), differentially expressed proteins were significantly downregulated at the P3 stage, while no significant alterations were revealed at later developmental stages (Fig. 2d).

**Table 1.**
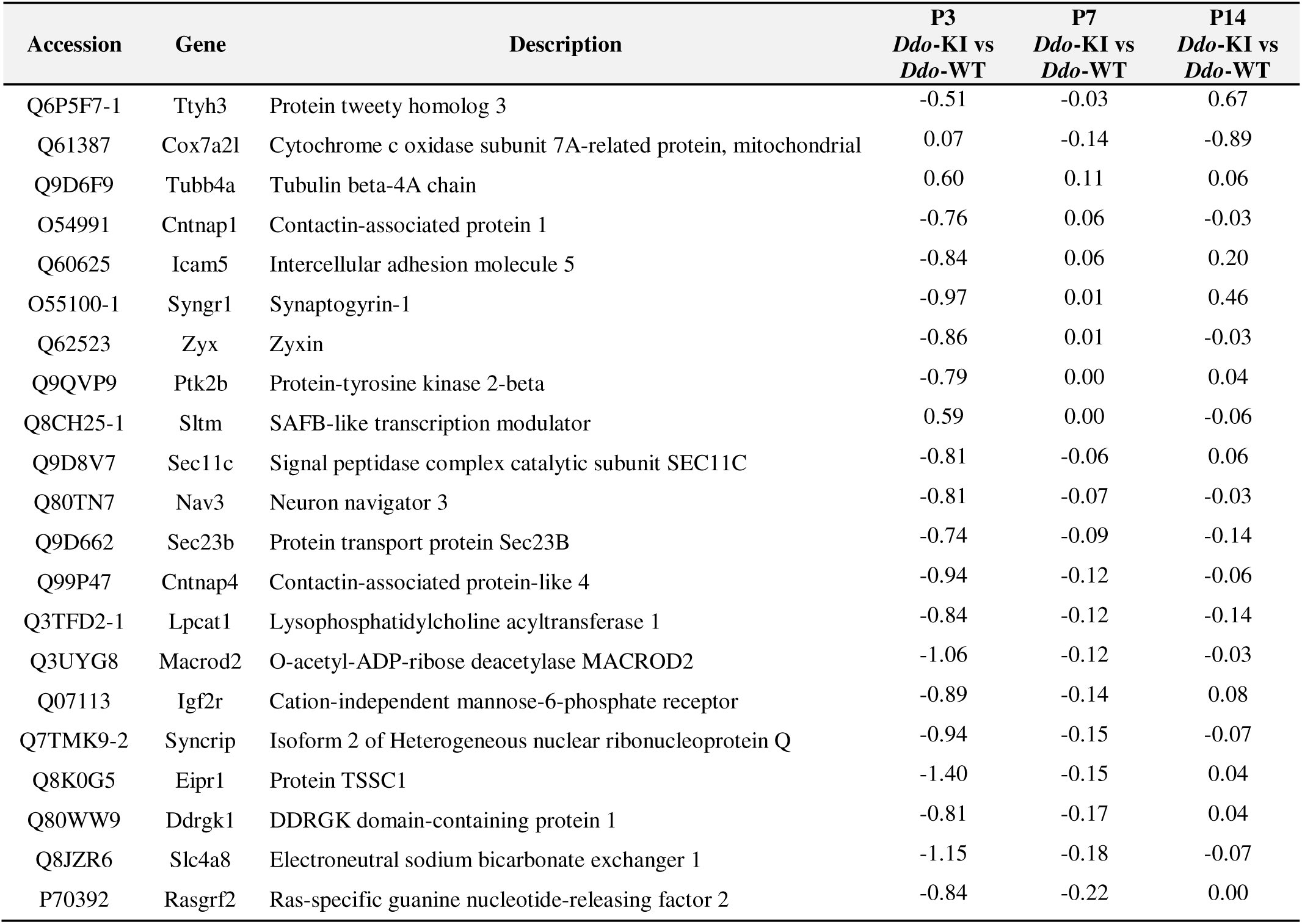
Differentially expressed proteins (0.66 ≥ FC ≥ 1.50) identified in the whole brain of *Ddo*-WT and *Ddo*-KI mice at postnatal day 3 (P3), P7, and P14 by quantitative nanoLC-MS/MS.

### D-aspartate depletion affects the expression of proteins involved in brain development, synaptic transmission and cytoskeletal organization

We next performed a gene ontology (GO) enrichment analysis on differentially expressed proteins, revealing that D-Asp-modulated proteins were mainly associated with molecular functions related to nervous system development, chemical synaptic transmission and its modulation, cellular component and cell junction organization, and cytoskeleton-dependent transport (Fig. 3a). The network of proteins involved in the most enriched affected functions is shown in Fig. 3b. Consistently, the most significantly enriched biological processes included synapse organization, axogenesis, cytoskeleton-dependent transport, transport along microtubules, regulation of neurotransmitter receptor activity and regulation of axogenesis (Fig. 4a). Notably, the most enriched cellular component categories were exclusively linked to synaptic structures (Fig. 4b). To further dissect the pathways influenced by D-Asp, we conducted a pathway enrichment analysis identifying the “Glutamatergic synapse” and the “Alanine, aspartate and glutamate metabolism” as the most significantly impacted pathways (Fig. 4c). Specifically, we mapped nine downregulated proteins within the “Glutamatergic synapse” pathway: Prkcg (↓0.43-fold), Grk2 (↓0.42-fold), Gria4 (↓0.48-fold), Homer1 (↓0.39-fold), Gria3 (↓0.41-fold), Adcy1 (↓0.70-fold), Slc17a7 (↓0.44-fold), Grin1 (↓0.42-fold) and Homer3 (↓0.72-fold). Additionally, three proteins involved in the “Alanine, aspartate and glutamate metabolism” pathway were significantly down-regulated: Gad1 (↓0.39-fold), Aldh4a1 (↓0.40-fold) and Adsl (↓0.43-fold). Overall, these findings highlight a regulatory role for D-Asp in modulating the expression of key proteins involved in glutamatergic neurotransmission.

**Figure 3.**
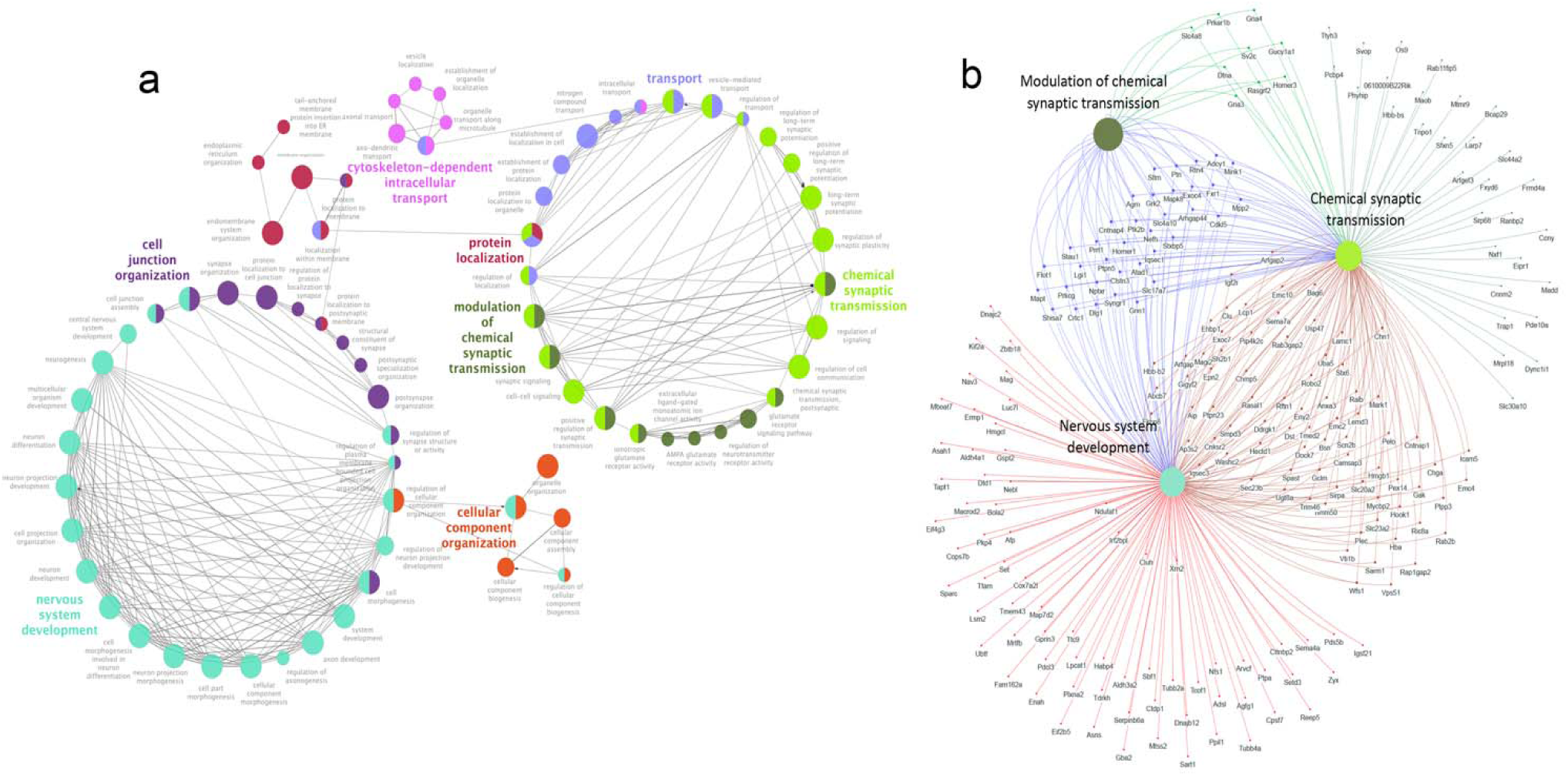
D-aspartate depletion affects the expression of proteins involved in brain development, synaptic transmission and cytoskeletal organization during early postnatal life. **(a)** Functionally grouped network of enriched molecular function categories for differentially expressed proteins in the whole brain of *Ddo*-KI and *Ddo*-WT mice at postnatal day (P) P3 and P14 generated using the ClueGO Cytoscape plug-in. The proportion of shared proteins between terms was evaluated using kappa statistics. GO terms are depicted as nodes whose size represents the term enrichment significance. **(b)** Venn network of proteins mapped within the top three enriched terms.

**Figure 4.**
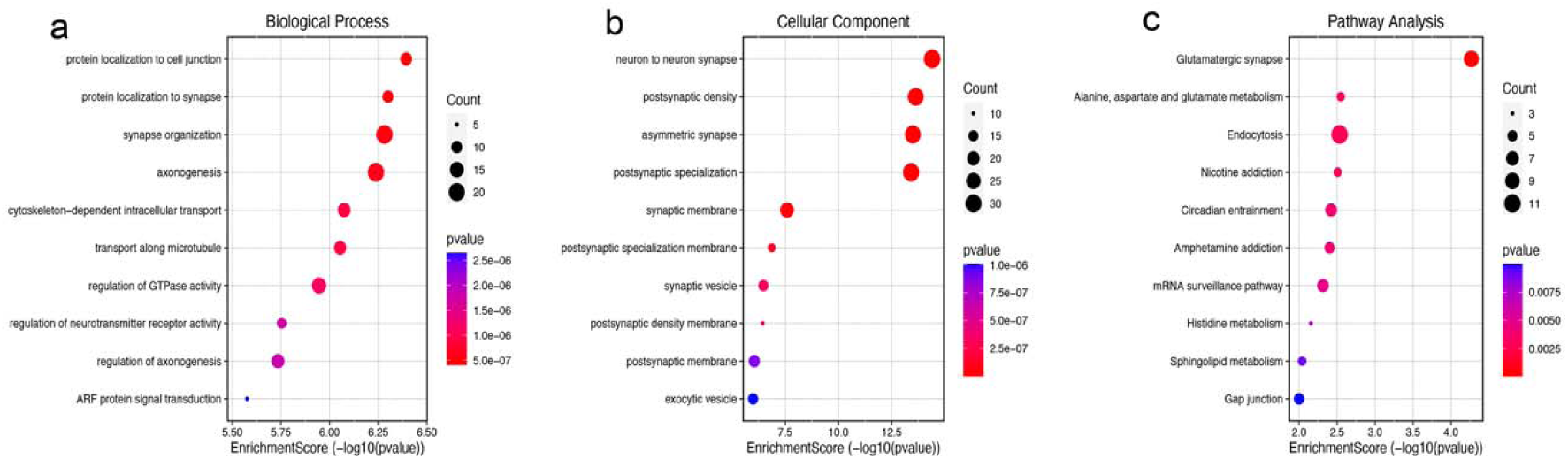
Analysis of enriched gene ontology terms at postnatal day 3. Enriched gene ontology (GO) terms for **(a)** “Biological process”, **(b)** “Cellular component” and **(c)** “Pathway analysis” of differentially expressed proteins in *Ddo*-KI and *Ddo*-WT mice at postnatal day 3. The colour and size of the plots represent the *p* value and the count of proteins, respectively.

### A subset of D-aspartate-modulated proteins in the early postnatal mouse brain is linked to autism spectrum disorder and schizophrenia

To gain insights into the potential involvement of D-Asp-modulated proteins in psychiatric disorders, we evaluated in the list of proteins affected by D-Asp depletion the prevalence of sequences previously related to autism spectrum disorders (ASD) and schizophrenia (SCZ). To this aim, we first extracted the subsets of *Homo sapiens* proteins associated with both psychiatric disorders by using the String: disease query plug-in of Cytoscape. Then, by using the g:Orth tool of the g:Profiler platform, we retrieved for these subsets the corresponding mouse orthologous protein entries that were subsequently matched against the list of D-Asp differentially expressed proteins. By this approach, we found that 15 differentially expressed proteins at the P3 developmental stage in *Ddo*-KI mice were previously associated with ASD (Fig. 5a, subsets A and B). GO enrichment analysis revealed that these proteins were mainly involved in functions including the regulation of synapse structure/activity, modulation of chemical synaptic transmission, nervous system development, glutamate receptor signalling pathway, behaviour and regulation of neurotransmitter levels (Fig. 5b; Suppl. Fig. 2a). Similarly, twenty-one differentially expressed proteins in *Ddo*-KI mice (20 at P3 and 1 at P14) were previously associated with SCZ (Fig. 5a, subsets A and C) and are involved in biological processes partially overlapping with those revealed for ASD (e.g. glutamate receptor signalling pathway and behaviour) as well as in the regulation of synaptic plasticity, glutamatergic synaptic transmission, postsynaptic chemical synaptic transmission, protein localization to synapse and regulation of axogenesis (Fig. 5c; Suppl. Fig. 2b). Interestingly, a comparison of D-Asp-modulated proteins with both psychiatric disorders revealed that nine proteins, i.e. Slc17a7 (↓0.44-fold), Gria3 (↓0.41-fold), Fxr1 (↓0.39-fold), Grin1 (↓0.42-fold), Mapt (↓0.46-fold), Homer1 (↓0.39-fold), Maob (↓0.44-fold), Cnnm2 (↓0.62-fold), Gad1 (↓0.39-fold), represent a common signature of potential relevance to study the role of D-Asp in neurodevelopmental disorders (Fig. 5a, subset A).

**Figure 5.**
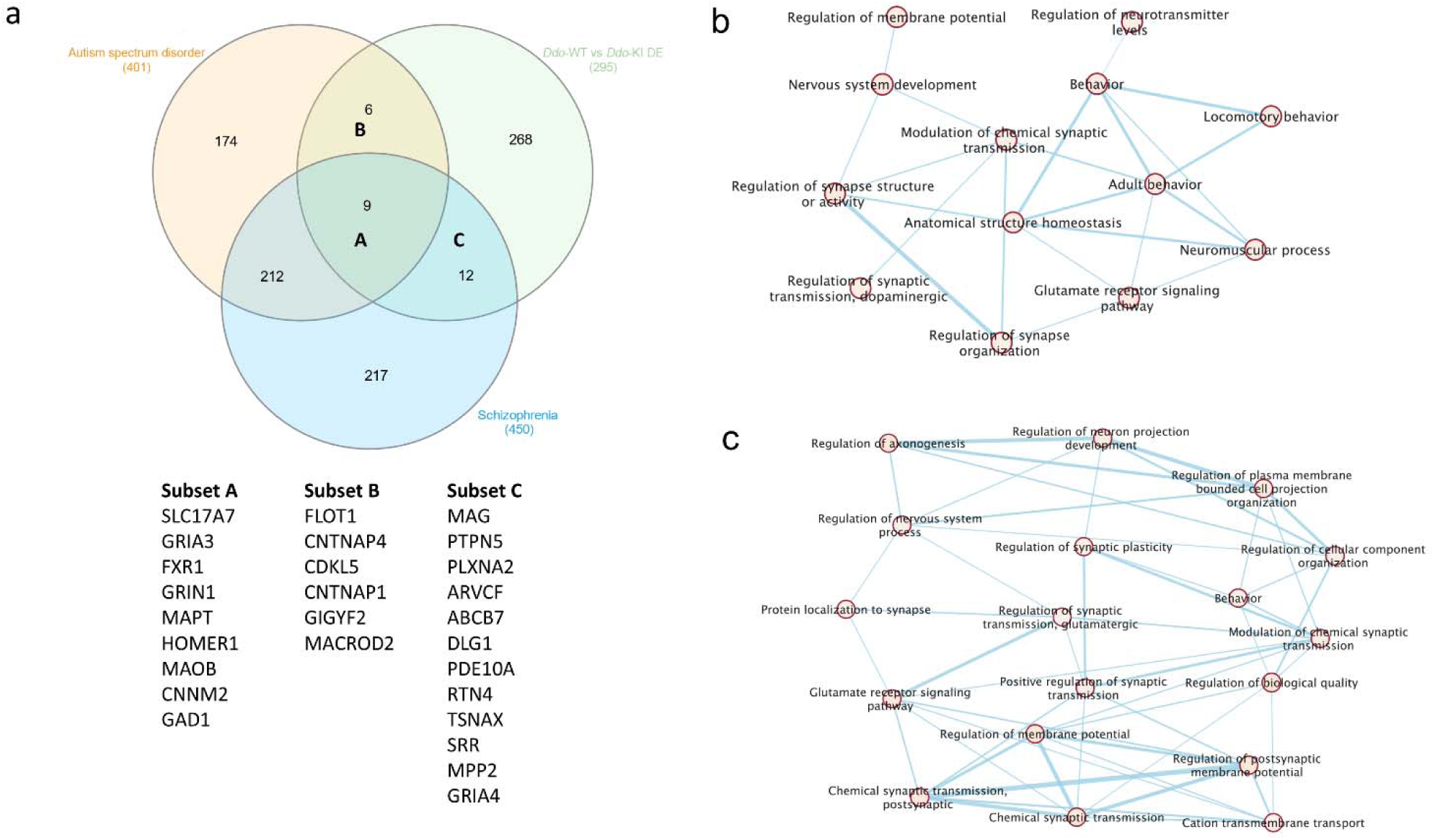
A subset of D-aspartate-modulated proteins in the whole brain of post-natal day 3 Ddo-KI mice is linked to autism spectrum disorder and schizophrenia. **(a)** Venn diagram showing the overlapping of proteins differentially expressed (DE) by D-aspartate in *Ddo*-WT vs *Ddo*-KI mice (*Ddo*-WT vs *Ddo*-KI DE) and orthologous proteins associated with autism spectrum disorder (ASD) and schizophrenia. **(b)** Enrichment map of biological processes involving proteins within the subsets A and B (*Ddo*-WT vs *Ddo*-KI DE and ASD) or **(c)** within the subsets A and C (*Ddo*-WT vs *Ddo*-KI DE and schizophrenia).

### Anticorrelation analysis of *DDO* in the human brain identifies genes overlapping with proteins downregulated in the *Ddo* knockin mouse brain at postnatal day 3

The above results and the previous finding of *DDO* gene duplication in a patient affected by thought disorders and ASD symptomatology^25^ suggest that *DDO* levels may influence the expression of genes critically implicated in this disorder, even in the human brain. To explore this possibility, we systematically analyzed the strongest positive and negative correlations between *DDO* and other genes in the human CNS, using a published tissue-specific coexpression network derived from a dataset of 544 human microarray experiments^27^. Among the 62 genes coexpressed with *DDO* (Suppl. Data 3) only one (*Pde10a*) overlapped with mouse proteins modulated in *Ddo*-KI mice. In contrast, the list of 126 *DDO*-anticorrelated genes showed nine sequences in common with mouse proteins modulated in *Ddo*-KI brains (*Fxyd6, Igsf21, Iqsec1, Madd, Nudt4, Oxr1, Pde1a, Ppm1h, Slc17a7*) (Suppl. Data 3), which represents a highly significant intersection (*p* = 0.00007, hypergeometric test). Notably, all the nine common proteins were downregulated in *Ddo*-KI transgenic brains at P3 (Suppl. Data 2). Interestingly, a GO enrichment analysis of *DDO*-anticorrelated genes in the human brain revealed a significant enrichment of terms associated with synaptic and cognitive functions.

### D-aspartate metabolism regulates the expression of *Grin1*, *Homer3*, *Syngr1* and *Iqsec1* genes in the early postnatal brain of *Ddo* knockin mice

We investigated whether the observed changes in protein expression may be a direct consequence of an altered gene expression caused by D-Asp depletion. To this aim, we performed qRT-PCR analysis on whole brain samples from *Ddo*-KI and *Ddo*-WT mice at P3 and P14. Specifically, we selected 30 genes encoding a subset of proteins identified in the mouse proteomic analysis, which are particularly relevant due to their role in glutamatergic transmission and/or brain development, as well as their known association with SCZ/ASD (Table 2; Suppl. Data 2). Some of these genes were also identified through *in silico* analysis based on anti-correlation/co-expression with *DDO* gene expression in the human CNS (Table 2; Suppl. Data 3). Consistent with proteomic data, statistical analysis revealed a significant reduction in *Grin1*, *Homer3*, *Syngr1* and *Iqsec1* mRNA expression in the brain of *Ddo*-KI mice at P3, compared with age-matched *Ddo*-WT control littermates (Table 2).

**Table 2.**
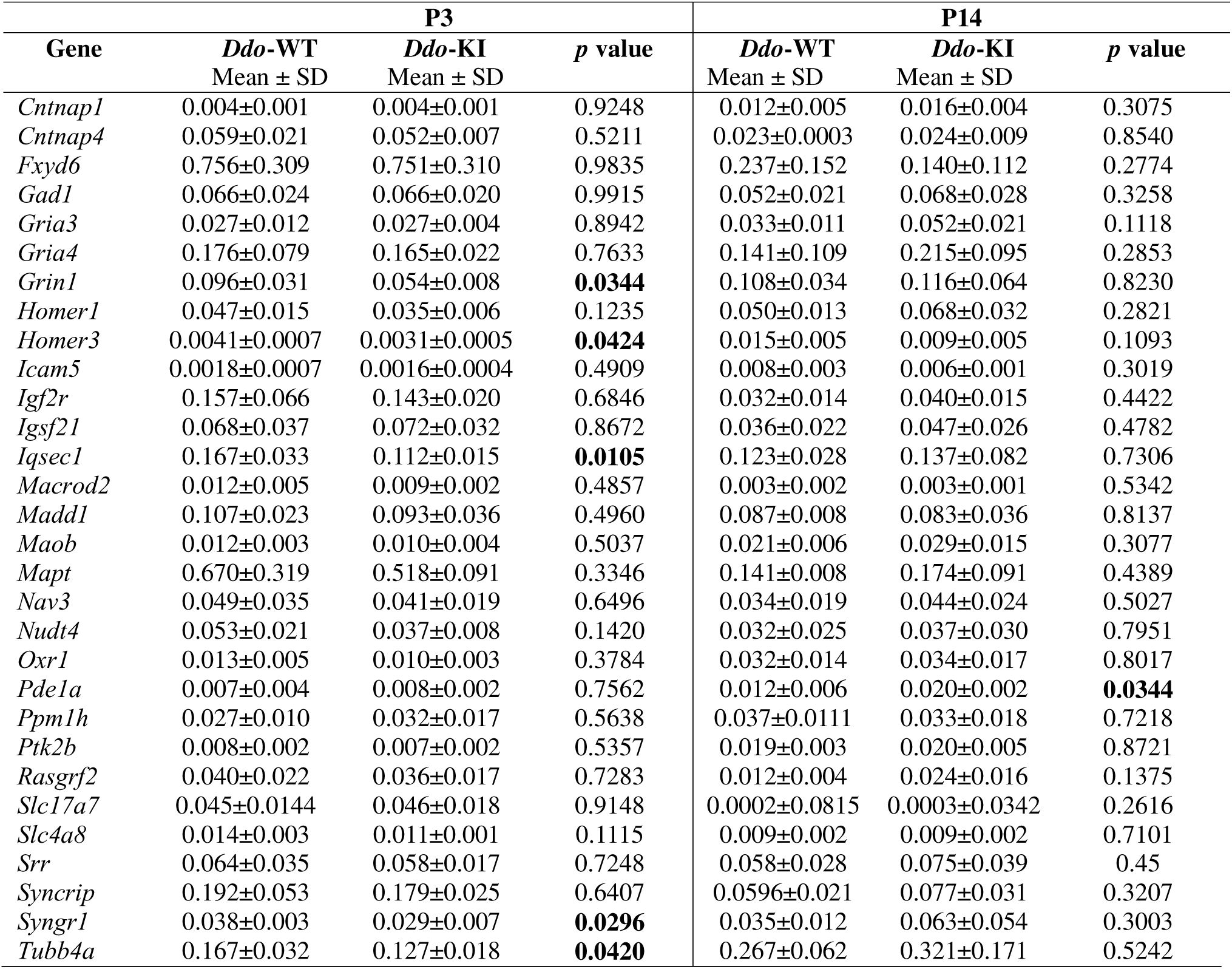
Analysis of mRNA expression of specific D-aspartate-modulated proteins in the whole brain of post-natal day 3 (P3) and P14 *Ddo*-KI and *Ddo*-WT mice by qRT-PCR.

Data are expressed as the difference in threshold cycle (2^-ΔΔCt^) between each specific gene and the reference gene, *Gapdh* (arbitrary units), and represented as the mean ± standard deviation (SD). Statistical analysis was performed by unpaired *t*-test. *p* < 0.05 was considered statistically significant and reported in bold.

## Discussion

Glutamatergic neurotransmission plays a crucial role in CNS development, regulating fundamental neurobiological processes such as neuronal proliferation, apoptosis, differentiation, migration and refinement of synaptic connections^28–32^. Accordingly, alterations in glutamatergic receptor function during prenatal and early postnatal development can lead to permanent changes in brain structure and organization, contributing to the later emergence of neurodevelopmental disorders such as SCZ and ASD^33–36^. Despite the influence of NMDAR and mGluR5 signaling in neurodevelopmental processes and related disorders, the precise contribution of diverse endogenous ligands and neurotransmitters regulating the activity of these glutamatergic receptors remains unclear.

Although research in this field is still limited, recent studies suggest that early depletion of cerebral D-Asp in *Ddo*-KI mice leads to morphological brain abnormalities, which in turn affect memory and social behaviour in juvenile and adult life^5, 25^, resembling ASD-like phenotypes reported in other animal models^37^. Similarly, another study demonstrated that prenatal and early postnatal supplementation of D-Asp in fragile X syndrome (*Fmr1* KO) mice improved fear conditioning memory and motor coordination deficit^38^. Further studies have shown that non-physiological elevation of this D-amino acid in adult mice improves cognitive functions^39^ and prevents SCZ-like phenotypes, including prepulse inhibition deficits, ataxia, and dysfunctional brain activation induced by the psychotomimetic NMDAR antagonists, dizocilpine (MK801) and phencyclidine (PCP)^18, 40, 41^.

In addition to preclinical investigations, independent *post-mortem* studies have shown a significant reduction in D-Asp levels in the PFC of SCZ patients compared to healthy controls^42, 43^. Consistently, a hypothesis-free machine learning algorithm identified a stable cluster of molecules, including the D-Asp/total Asp ratio, in the *post-mortem* dorsolateral PFC (DLPFC) effectively distinguishing SCZ patients from non-psychiatric controls^44^. In line with *post-mortem* studies, recent findings have revealed dysregulated D-Asp concentrations in the blood samples from living SCZ patients^45, 46^. Moreover, the identification of a chromosome 6 duplication, including the entire *DDO* gene, in a young patient with severe intellectual disability, thought disorders and behavioural abnormalities reminiscent of ASD and SCZ symptomatology^25^ provides further genetic evidence linking D-Asp metabolism to neurodevelopmental disorders. Despite these advances, whether D-Asp metabolism influences NMDAR and mGluR5 transmission through selective protein expression patterns remains unclear.

In this study, we found that D-Asp depletion significantly alters the proteome profile of the developing mouse brain at an early postnatal stage. Notably, we revealed that in *Ddo*-KI mice most of the differentially expressed proteins (cut-off level > 1.3-fold) were detected at P3, suggesting that the regulatory effect of D-Asp on protein expression is confined in a narrow postnatal developmental window.

GO enrichment analysis revealed that proteins modulated by D-Asp are primarily associated with nervous system development, chemical synaptic transmission and its modulation, and cytoskeleton-mediated intracellular transport, confirming previous *in vivo* and *ex vivo* studies^5, 15, 16, 18–22, 24, 25, 38, 39^

Among the most significantly altered proteins (cut-off > 1.3-fold), we identified key components of the glutamatergic synapse affected by D-Asp depletion, including the essential NMDAR subunit, GluN1 (encoded by the *Grin1* gene), the AMPAR subunits, GluA3 (*Gria3*) and GluA4 (*Gria4*), and the post-synaptic density proteins, Homer1 and Homer3^47^. Moreover, we identified GAD1, an enzyme central to GABA synthesis, and thus indirectly associated with glutamatergic neurotransmission^48^. Notably, a previous study has revealed an altered number of parvalbumin-positive GABAergic interneurons in the cortex of *Ddo*-KI mice^5^, supporting a potential implication of D-Asp in regulating inhibitory circuits.

Applying a more stringent protein expression cut-off (> 1.5-fold), we further refined a subset of proteins involved in nervous system development and chemical synaptic transmission. Many of these proteins directly or indirectly interact with NMDARs and/or mGluR5, contributing to the modulation of glutamate-mediated synaptic activity. Notable examples include CNTNAP1, which regulates NMDAR-dependent synaptic plasticity^49^ and mGluR5-dependent hippocampal memory formation^50^, and ICAM5, which modulates dendritic outgrowth and spine maturation by competing with NMDARs^51–53^. Additionally, SYNGR1 is involved in synaptic vesicle function and significantly impacts NMDAR-dependent long-term potentiation (LTP)^54^. Moreover, mGluR5-mediated induction of PTK2B (also known as PYK2) promotes NMDAR phosphorylation, thereby contributing to intracellular Ca^2+^ homeostasis^55, 56^. Finally, RASGRF2 is essential for NMDAR-dependent hippocampal LTP and spine enlargement^57, 58^, while SLC4A8 modulates the pH-dependent release of neuronal glutamate^59^.

Among D-Asp-regulated proteins, several play a key role in processes critical for brain development, including synapse formation, synaptic plasticity and neurotransmission (SYNGR1, CNTNAP4, ICAM5, PTK2B, RASGRF2, GAD1)^54, 60–64^, neuronal migration and axon guidance (CNTNAP1, TUBB4A, ICAM5, NAV3)^60, 65–67^, myelination (TUBB4A, CNTNAP1)^67, 68^, regulation of signaling pathways (IGF2R, SLC4A8)^69, 70^ and gene expression (SYNCRIP)^71^, cytoskeletal organization, dendrite morphogenesis and structural integrity (MAPT, TUBB4A, PTK2B)^65, 72, 73^.

D-Asp has been shown to induce NMDAR-dependent late-phase LTP and dendritic spinogenesis in the rodent hippocampus by modulating actin polymerization mechanisms^20, 24^. Consistently, findings in the mouse testis indicate that D-Asp influences spermatogenesis by regulating actin- and tubulin-mediated cytoskeleton dynamics^74, 75^. The present results confirm the impact of D-Asp on cerebral protein levels implicated in cytoskeleton-dependent transport and remodeling, including microtubule binding and stability (MAPT, TUBB4A)^65, 73^, actin regulation (PTK2B, RASGRF2)^64, 73^, cytoskeleton adhesion (CNTNAP1, ICAM5)^60, 67^, and microtubule organization and axon guidance (NAV3)^66^.

In line with the increasing evidence linking early cerebral D-Asp to neurodevelopmental psychiatric disorders, our proteomic analysis identified a subset of D-Asp-regulated proteins at P3 that are associated with ASD, SCZ or both disorders. Many of these proteins are known to modulate glutamatergic neurotransmission. Among them, we identified the aforementioned GluN1, GluA3, GluA4, Homer1, the synaptic vesicle-associated glutamate transporter, SLC17A7/VGLUT1^76^, the biosynthetic enzyme of glutamate-GABA conversion, GAD1^48^, as well as CNTNAP1^49^, FXR1^77^, FLOT1^78^, CDKL5^79^, PTPN5^80^, DLG1/SAP97^81^, RTN4/NogoA^82^ and MPP2^83^.

To further support our proteomic findings, we analyzed publicly available human datasets for genes most strongly correlated with *DDO* expression. Among them, we identified nine anti-correlated human genes encoding orthologous proteins to those found in proteomic analysis. Beyond *Slc17a7*, which directly impacts glutamatergic transmission^76^, the majority of these genes encode proteins that modulate glutamatergic functions, affecting synaptic transmission (*Iqsec1, Ppm1h, Pde1a*)^84–86^, neuronal excitability (*Fxyd6, Madd*)^87, 88^, synapse organization (*Igsf21*)^89^ or neuroprotection (*Oxr1*)^90^. Many of these also play a critical role in brain development (*Slc17a7, Iqsec1, Ppm1h*, *Fxyd6*)^76, 84, 85, 87^ and have been previously linked to neurodevelopmental psychiatric disorders (*Slc17a7, Iqsec1, Pde1a, Madd*, *Fxyd6*)^91–95^.

To investigate whether the modulatory effect of D-Asp on protein expression is somehow reflected at the mRNA level, we performed qRT-PCR analysis in the brain of *Ddo*-KI mice at P3 and P14. We focused on genes encoding a subset of differentially expressed proteins relevant to glutamatergic transmission, brain development, and/or neurodevelopmental psychiatric disorders. Some of these genes were also strongly anti-correlated with *DDO* expression in human *in silico* database. Among the tested genes, we found that *Grin1*, *Homer3*, *Syngr1* and *Iqsec1* were significantly downregulated in *Ddo*-KI mice compared to *Ddo*-WT controls at P3 but not at P14, in line with a selective, time-dependent decrease in the corresponding protein levels. Regardless of the precise molecular mechanism involved, our present data highlight that D-Asp metabolism physiologically influences the brain proteome at P3 affecting both transcriptional and translational processes.

Although further targeted studies are needed, we hypothesize that the selective impact of D-Asp metabolism on the brain proteome during the early postnatal stage aligns with a critical role of NMDAR signaling in regulating neurodevelopmental processes relevant to psychiatric disorders^96, 97^. Specifically, the narrow time window during which D-Asp affects brain proteome may coincide with the developmental shift from GluN2B to GluN2A subunit expression of NMDARs^98^. This transition is crucial for the maturation of synaptic signaling and cognitive functions, and alterations in its timing could influence neurodevelopmental pathways implicated in psychiatric conditions^98^.

In conclusion, the present findings highlight that D-Asp modulates the expression of key proteins implicated in NMDAR and mGluR5-dependent processes that are crucial for neurodevelopment and associated neuropsychiatric disorders. These observations emphasize the need for further research to elucidate the molecular mechanisms underlying D-Asp signaling in neurodevelopment and highlight the translational potential of D-Asp as a target for future studies exploring pathophysiological mechanisms of these disorders and their related therapeutic interventions.

## Methods

### Animals

*Ddo*-KI and *Ddo*-WT mice were generated and genotyped by PCR according to the protocol previously reported^5^. Animals were housed in groups (4–5 per cage), at a constant temperature (22 ± 1 °C) on a 12 h light/dark cycle (lights on at 7 AM) with food and water ad libitum. All research involving animals was carried out following the directive of the Italian Ministry of Health governing animal welfare and protection (D.LGS 26/2014) and approved by “Direzione Generale della Sanità e dei Farmaci Veterinari (Ufficio 6)” (permission nr. 796/2018).

### HPLC analysis

The levels of D-Asp, L-aspartate, L-glutamate, L-asparagine, D-serine, L-serine, glycine, L-threonine, L-arginine, taurine, L-alanine, and GABA in mouse brain tissues were analysed by HPLC. Brain samples were pulverized in liquid nitrogen and an aliquot of each (7-18 mg) was homogenized in 1:10 (w/v) 0.2 M trichloroacetic acid (TCA), sonicated (4 cycles, 10 s each), and centrifuged at 13 000 ×g for 20 min. TCA supernatants containing amino acids were then neutralized with NaOH and subjected to pre-column derivatization with o-phthaldialdehyde (OPA)/N-acetyl-L-cysteine (NAC). Diastereoisomer derivatives were resolved on a UHPLC Agilent 1290 Infinity (Agilent Technologies, Santa Clara, CA, USA) using a ZORBAX Eclipse Plus C8, 4.6 × 150 mm, 5 μm (Agilent Technologies, Santa Clara, CA, USA) under isocratic conditions (0.1 M sodium acetate buffer, pH 6.2, 1% tetrahydrofuran, and 1.5 mL/min flow rate). Each run was followed by a washing step with 0.1 M sodium acetate buffer, 3% tetrahydrofuran, and 47% acetonitrile. The precipitated protein pellets were solubilized in 1% SDS solution and quantified by bicinchoninic acid (BCA) assay method (Pierce™ BCA Protein Assay Kits, (Thermofisher scientific, Rockford, IL, USA). Identification of amino acids was based on retention time, compared with the one associated with external standards. Each compound was quantified using 6-15 point calibration curve generated using external standards in the range of the predicted analyte concentration in the sample. The detected amino acid concentration in tissue homogenates was normalized by the total protein content and expressed as nmol/mg protein. If the amino acid concentration in the tissue analysed was below the limit of detection (0.01 pmol) the value was set at 0.

### Quantitative RT-PCR

The expression profile of *Ddo* and 30 other genes, including *Gria3*, *Syncrip*, *Icam5*, *Tubb4a*, *Cntnap1*, *Ptk2b*, *Maob*, *Slc17a7*, *Pde1a*, *Macrod2*, *Homer1*, *Slc4a8*, *Mapt*, *Rasgrf2*, *Grin1*, *Igf2r*, *Gad1*, *Gria4*, *Cntnap4*, *Homer3*, *Iqsec1*, *Madd*, *Nav3*, *Nudt4*, *Oxr1*, *Srr*, *Syngr1*, *Ppm1h*, *Igsf21*, *Fxyd6*, was evaluated on total RNAs of *Ddo*-KI and *Ddo*-WT (n = 5 mice/genotype/age). RNA was isolated by using TRIzol RNA Isolation Reagents (Thermo Fisher Scientific, Waltham, MA, USA) protocol. Five μg of total RNA were reverse transcribed with the Superscript III-First strand kit (Thermo Fisher Scientific, Waltham, MA, USA). qRT-PCR reactions were performed in triplicate, using gene-specific primers (Suppl. Table 1) and ITaq Universal Sybr Green Supermix (Bio-Rad, Hercules, CA, USA) following the manufacturer’s directions. Results were normalized versus the expression of the glyceraldehydes-3-phosphate dehydrogenase (*Gapdh*) gene. Data were analyzed with unpaired *t*-test to compare the gene expression between genotypes at each age. A value of *p* < 0.05 was considered to be statistically significant.

### Sample preparation for proteomic analyses

Proteomic analyses were performed on pooled (n = 3) whole brains of *Ddo*-KI and *Ddo*-WT mice and collected at different stages of postnatal development: P3, P7 and P14. Animals were sacrificed and the whole brains were dissected within 20 s on an ice-cold surface. All tissue samples were pulverized in liquid nitrogen and stored at −80 °C for subsequent processing. Tissues were then lysed in ice-cold lysis buffer (100 mM Triethylammonium bicarbonate TEAB, SDS 1%) and disrupted by two cycles of sonication at a 20% amplitude for 30 s on ice. Lysates were cleared by centrifugation at 16,000 × g for 15 min at 4°C. Supernatants were transferred into new tubes, treated with 1 U of RQ1 Dnase (Promega) for 30 min at room temperature, and the protein concentration was determined by using the Pierce BCA Protein assay kit (Thermo Scientific, Rockford, IL). For each condition, equal amounts of proteins (50 µg in 100 µL of 100 mM TEAB) were reduced with 10 mM Tris-(2-carboxyethyl)-phosphine (TCEP) for 1 h at 55 °C and alkylated with 18 mM iodoacetamide by incubating samples for 30 min at room temperature in the dark. Proteins were then precipitated overnight by adding six volumes of pre-chilled acetone. Following centrifugation at 8,000 × g for 10 min at 4°C, protein pellets were resuspended in 100 µL of 100 mM TEAB and digested overnight with MS grade trypsin (Pierce) at an enzyme/substrate ratio of 1:40 at 37°C. The resulting peptide mixtures were chemically labelled with the TMT isobaric tags as previously reported^99, 100^ by using the following isobaric tags: 131 P3 WT, 130C P3 KI, 129N P7 WT, 128C P7 KI, 127N P14 WT, 126 P14 KI. Briefly, 0.8 mg of TMT reagents in 41 µL of anhydrous acetonitrile were added to each sample. The reaction proceeded for 1 h and then was quenched for 15 min with hydroxylamine to a final concentration of 0.3%. Two mixtures of samples were then prepared by combining the six conditions for both embryonic and postnatal samples at equal amounts. Mixed samples were diluted in 0.1% TFA/2% CH_3_CN to a final concentration of 0.5 µg/µL for LC-MS analyses.

### High-resolution nanoLC−Tandem Mass Spectrometry

Aliquots of TMT labelled samples (2.5 µg) were analyzed by high-resolution nanoLC-Tandem Mass Spectrometry using a Q-Exactive Orbitrap mass spectrometer equipped with an EASY-Spray nano-electrospray ion source (Thermo Fisher Scientific, Bremen, Germany) and coupled to a Thermo Scientific Dionex UltiMate 3000RSLC nano system (Thermo Fisher Scientific)^26^. The solvent composition was 0.1% formic acid in water (solvent A) and 0.1% formic acid in acetonitrile (solvent B). Peptides were loaded on a trapping PepMap™100 μCartridge Column C18 (300 μm x 0.5 cm, 5 μm, 100 Å) and desalted with solvent A for 3 min at a flow rate of 10 μL/min. After trapping, eluted peptides were separated on an EASY-Spray analytical column (50 cm x 75 μm ID PepMap RSLC C18, 3 μm, 100 Angstrom), heated to 35 °C, at a flow rate of 300 nL/min applying the following gradient: 5% B for 3 min, from 5% to 27.5% B in 222 min, from 27.5% to 40% B in 10 min, from 40% to 95% B in 1 min. Washing (95% B for 4 min) and re-equilibration (5% B for 24 min) steps were always included at the end of the gradient. Eluting peptides were analysed on the Q-Exactive mass spectrometer operating in positive polarity mode with a capillary temperature of 280 °C and a potential of 1.9 kV applied to the capillary probe^100, 101^. Full MS survey scan resolution was set to 70000 with an automatic gain control (AGC) target value of 3 × 10^6^ for a scan range of 375-1500 m/z and maximum ion injection time (IT) of 60 ms. The mass (m/z) 445.12003 was used as lock mass. A data-dependent top 12 method was operated during which high-energy collisional dissociation (HCD) spectra were obtained at 35000 MS2 resolution with AGC target of 1 × 10^5^ for a scan range of 200-2000 m/z, maximum IT of 120 ms, 1.6 m/z isolation width and normalized collisional energy (NCE) of 32. Precursor ions targeted for HCD were dynamically excluded for 30 s. Full scans and Orbitrap MS/MS scans were acquired in profile mode, whereas ion trap mass spectra were acquired in centroid mode. Charge state recognition was enabled by excluding unassigned and 1, 7, 8, >8 charged states. All data were acquired with Xcalibur 3.1 software (Thermo-Fisher Scientific).

### Protein identification and quantitation

For data processing, the acquired raw files were analyzed with the Thermo Scientific Proteome Discoverer 2.4 software (Thermo Fisher Scientific) using the SEQUEST HT search engine. The HCD MS/MS spectra were searched against the *Mus musculus* database (UniProt release 2023_01, number of entries 17761 sequences) assuming trypsin (Full) as digestion enzyme and two allowed number of missed cleavage sites. Mass tolerances were set to 10 ppm and 0.02 Da for precursor and fragment ions, respectively. The oxidation of methionine (+15.995 Da) was set as a dynamic modification. Carbamidomethylation of cysteine (+57.021 Da) and the TMT label on lysines and the N-terminus (229.1629) were set as static modifications. False discovery rates (FDRs) for peptide spectral matches (PSMs) were calculated and filtered using the Percolator node in Proteome Discoverer that was run with the following settings: Maximum Delta Cn 0.05, a strict target FDR of 0.01, a relaxed target FDR of 0.05 and validation based on q-value. Protein identifications were accepted when the protein FDR was below 1 % and when present in at least two out of three replicate injections with at least two peptides.

### Bioinformatic analyses

The clustered heatmap of the proteins that were differentially expressed with log2 fold change values - 0.6 ≥(log2fold change)≥ 0.6 in at least one out of three replicates was generated by hierarchical cluster analysis (Euclidean average distance) by using the Perseus software platform (version 2.0.11) after Z-score normalization of the protein dataset, and minimum and maximum intensity values were represented by default green and red colours, respectively. For bioinformatic analyses, proteins with log2 fold change values -0.4 ≥ (log2fold change) ≥ 0.4 were considered as differentially expressed (DE). Molecular function enrichment analysis was performed by using the ClueGO plug-in of Cytoscape 3.6.0 to generate a functionally grouped GO/pathway term network of enriched molecular function categories for the identified proteins based on kappa statistics^102, 103^. For further data visualization, the results of this analysis were imported onto the Evenn platform to generate the Venn network of proteins mapped within enriched terms^104^. The enriched Gene Ontology (GO) terms for DE proteins were extracted by using a module integrating the clusterProfiler R package^105^. Enrichment plots and GOChord plots were generated and visualized by using the online platform for data analysis and visualization available at https://www.bioinformatics.com.cn. The networks of *Homo sapiens* proteins associated with autism spectrum disorders and schizophrenia were retrieved by using the String: disease query plug-in of Cytoscape using a confidence (score) cutoff of 0.4. The g:Orth tool of the g:Profiler web service platform was then used to retrieve mouse orthologous protein mappings of *Homo sapiens* entries based on the gene information from the Ensembl database^106^. This subset of disease-related proteins was subsequently matched against the list of D-Asp differentially expressed proteins to map modulated proteins associated with the selected neurodegenerative disorders.

The lists of human genes showing strong coexpression or anticorrelation with DDO was obtained from the dataset previously described^27^, using as coexpression or anticorrelation cutoff the presence in top or bottom 1% ranks of the Pearson correlation coefficients, respectively, with reciprocity criterion. The Gene Ontology enrichment analysis of these lists was performed using the DAVID analysis tool^107^ and the statistical significance of intersections was assessed using the Fisher exact test.

## Acknowledgments

We kindly thank Mariangela Valletta for her support in mass spectrometry analyses and Arianna De Rosa for her support in mouse genotyping and critical discussion.

## Author Contributions

F.E. supervised HPLC experiments, wrote and edited the paper. R.R. performed proteomic experiments, acquired data, prepared the figures and wrote the paper. F.C. performed qRT-PCR experiments and acquired data. T.N. supervised HPLC experiments. R.d.V. performed HPLC experiments, acquired data and prepared the figures. E.C. performed proteomic experiments. P.V.P. edited the paper. F.D.C. performed analysis on human *in silico* database and edited the paper. T.E. supervised qRT-PCR experiments and edited the paper. A.U. conceived and designed the study, and edited the paper. A.C. supervised proteomic experiments and edited the paper.

## Data and materials availability

All data needed to evaluate the conclusions of the work are present in the paper and the supplementary materials.

