## Supplementary information for "Tandem Mass Tag-Based High-Resolution LC-MS/MS identifies free D-aspartate-induced expression of proteins linked to schizophrenia and autism spectrum disorder"

**^@^ Corresponding authors:**

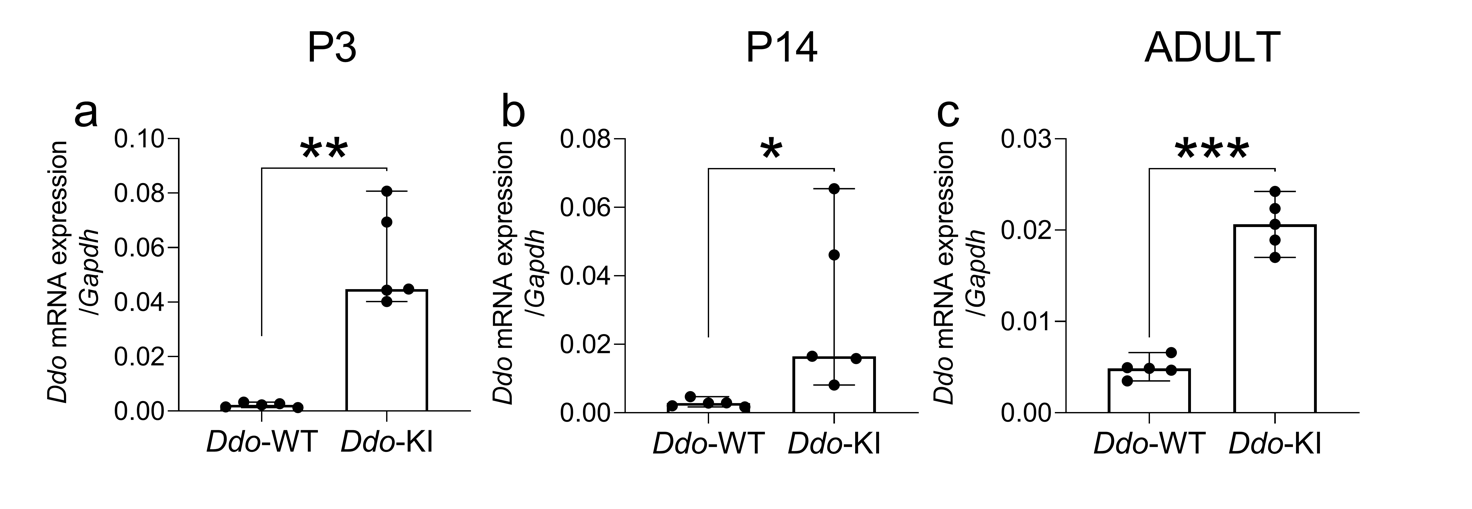

**Supplementary Figure 1.** *Ddo mRNA expression levels in the whole brain of Ddo-KI and Ddo-WT mice during postnatal ontogenesis.* Analysis of *Ddo* mRNA levels in the whole brain of *Ddo*-KI and *Ddo*-WT mice at **(a)** postnatal day 3 (P3) **(b)** P14 and **(c)** adult phase (at approximately P300). Data are expressed as the difference in threshold cycle (2^-ΔΔCt^) between *Ddo* gene and the reference gene, *Gapdh* (arbitrary units), and shown as box and whisker plots representing the median with interquartile range (IQR). Dots represent individual mice values. **p* < 0.05, ***p* < 0.01, ****p* < 0.0001, compared with age-matched *Ddo*-WT mice (unpaired *t*-test).

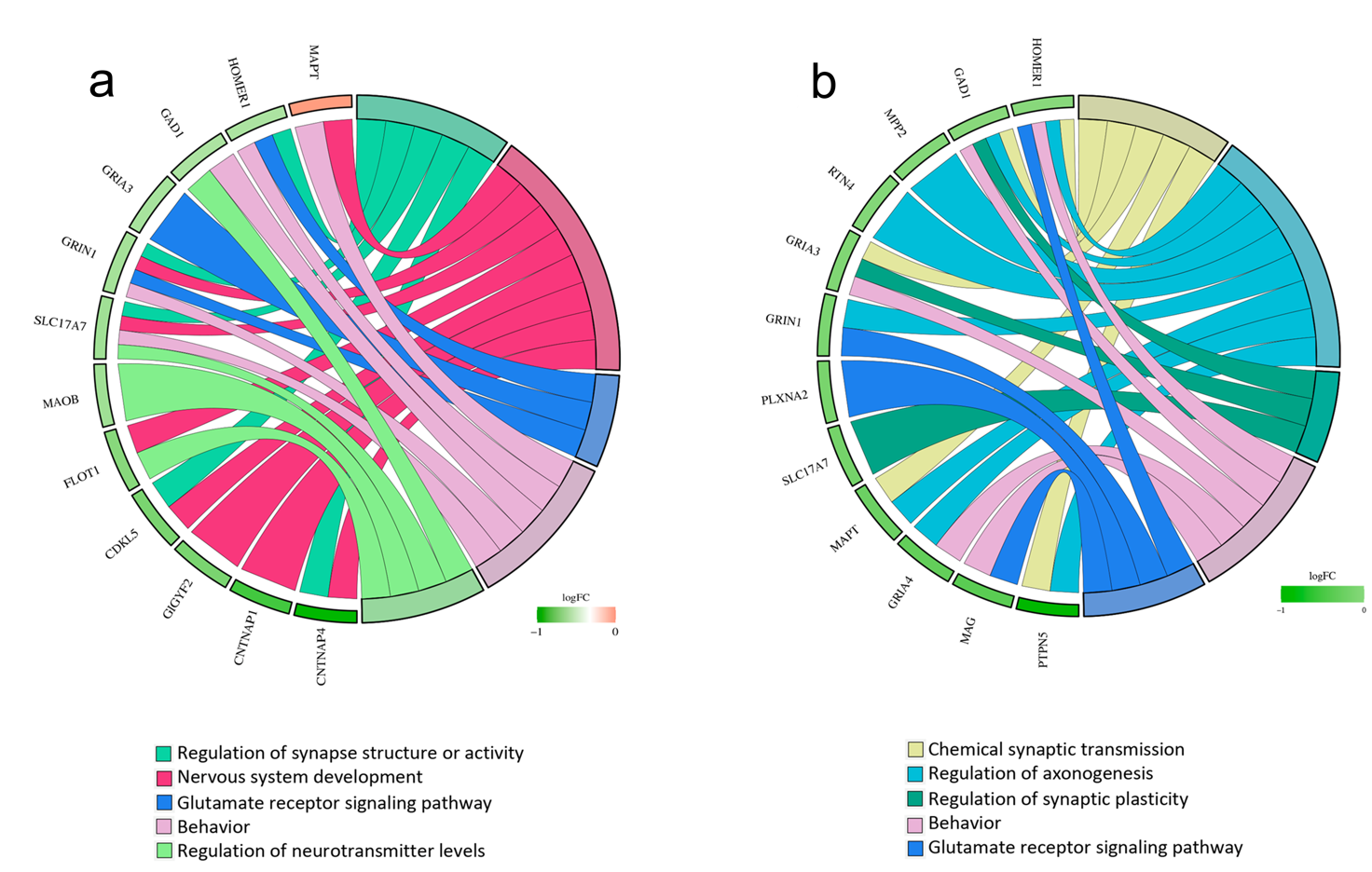

**Supplementary Figure 2.** *Key brain processes modulated by D-aspartate-regulated proteins associated with autism spectrum disorder or schizophrenia.* Gene Ontology (GO) chord plots showing relationships between selected enriched biological process GO terms and D-aspartate-regulated proteins overlapping with **(a)** autism spectrum disorders and **(b)** schizophrenia.

| **Gene** | **Primer forward (5’-3’)** | **Primer reverse (5’-3’)** | **Reference** | **Product size** | **MT** |
| --- | --- | --- | --- | --- | --- |
| *Cntnap1* | CACTCCAACCAGACAGCATTCCA | CGGTAAGTTCGCAGTAGCAGATGA | 1 | 220 | 60°C |
| *Cntnap4* | TTTGGAACGCAGCTTCCTTTA | GAGAGGGCTGTCGTCTTGAAA | 2 | 109 | 60°C |
| *Ddo* | ACCACCAGTAATGTAGCGGC | GGTACCGGGGTATCTGCA | 3 | 62 | 60°C |
| *Fxyd6* | TTGGTGTTTGCTGTGGTCCT | TCCGCAGCGTTTGTAGTGAT | This paper | 143 | 60°C |
| *Gad1* | TCCTGGTTGACTGTAGAGACAC | CATATTGGTATTGGCAGTCGAT | 4 | 137 | 60°C |
| *Gapdh* | GGTGCTGAGTATGTCGTGGA | GTGGTTCACACCCATCACAA | 5 | 141 | 60°C |
| *Gria3* | ACAAAGCCCTCCTGATCCTC | GTGAAGAACCACCAAACCCC | 5 | 144 | 60°C |
| *Gria4* | TGTTGGGAAGCACGTCAAAG | TCGTCACCATGGGCGTATTA | 5 | 140 | 60°C |
| *Grin1* | GTTCTTCCGCTCAGGCTTTG | AGGGAAACGTTCTGCTTCCA | 6 | 66 | 60°C |
| *Homer1* | GCAAAGGAGAAGTCGCAGG | CCTTGGCTCTGAGTTCTGT | This paper | 153 | 60°C |
| *Homer3* | CAGACAGTCGAGCCAACACT | ATTTCTCTCGAGCCAGCCTG | This paper | 117 | 60°C |
| *Icam5* | AGAACAGGAAGGCACCAAACAG | CTGGCTCACTCAAAGTCAGAAGAG | 7 | 122 | 60°C |
| *Igf2r* | TCAGTTTGTCTGTTCGGACCG | TAGACACGTCCCTCTCGGACTT | 8 | 164 | 60°C |
| *Igsf21* | TGCGTGAGATCGTGTGGTAC | GCGGTAATTCTCCATGTGCG | This paper | 110 | 64.3 |
| *Iqsec1* | GGTGTGGATGACGGTGAAGA | TTCTTGCCCACGATGAGCTT | This paper | 137 | 60°C |
| *Macrod2* | GCCTGAGACGGTTATGGAAA | TGTCTCCCACCCTTCTTGTC | 9 | 212 | 60°C |
| *Madd* | TACATGTTCCCGGTCATCCC | GCAATCACCCTATTGCTGTC | This paper | 176 | 60°C |
| *Maob* | TCCATGGAAAGCACCCCTTG | GAGCGTGGCAATCTGCTTTG | This paper | 103 | 60°C |
| *Mapt* | GGGTCAGGTGAACCACCAAA | GTGTTGGTAGGGATGGGGTG | This paper | 103 | 60°C |
| *Nav3* | GGTCTCCAGCACATCATCC | GTGAGCATTTGCTGACAGCT | This paper | 139 | 60°C |
| *Nudt4* | GGTGCTGCTGTGAGGGAAG | ATGTTCTGTGCTTCCGGTCT | This paper | 103 | 60°C |
| *Oxr1* | TGAAGAACTGCGCACACTCT | ACTGGGGTCGCTTAGGTTTG | This paper | 130 | 60°C |
| *Pde1a* | GAAGCAAGCGGGGAGCATAG | AAAGGCAATTAGGCAAGAAACAGG | 10 | 105 | 60°C |
| *Ppm1h* | GCCCCATCCTCATCCTCAAG | TGGAGTTTCTGTTCGGGGTG | This paper | 168 | 60°C |
| *Ptk2b* | CAGATGACCGTGGGCGAAGT | GGCAAGTAGCGGATTTGAAGG | 11 | 89 | 60°C |
| *Rasgrf2* | GTGAGGGCCAGAAAGCTGTCTTTGACGTCT | TCGGCTACCTGTCCTCCAGGC | 12 | 250 | 60°C |
| *Slc17a7* | CTGGGGTCCTTGTGCAGTAT | GTATTTGCGCTCCTCCTCAG | 5 | 146 | 60°C |
| *Slc4a8* | GGGCAGCAGTACCATGAGAT | GTCCAGGAACTCGTCAATCC | 13 | 126 | 60°C |
| *Srr* | CCCTTGGTAGATGCACTGGT | TCAGCAGCGTATACCTTCACA | 3 | 107 | 60°C |
| *Syncrip* | ACCTTGCCAACACGGTAACA | CCATAGCCTTGACAGCACCA | 14 | 135 | 60°C |
| *Syngr1* | CGTCGTGTCTTGGGTCTTCT | TAGAGCAGGCAGGTGAGGAA | This paper | 168 | 60°C |
| *Tubb4a* | CCAGATCTTTCGGCCAGACA | GGCATCCACTAACTCCGCG | This paper | 100 | 60°C |

**Supplementary Table 1.** The complete list of gene-specific primers used for qRT-PCR.

MT: melting temperature.

**References**
